# Immediate transcriptional changes initiated by direct cell-cell contact between cytotoxic T cells and cancer cells

**DOI:** 10.64898/2026.02.17.706359

**Authors:** Zhouhui Qi, Madeleine Lehander, Narmadha Subramanian, Eirini Giannakopoulou, Johanna Olweus, Michael Hagemann-Jensen, Christoph Ziegenhain, Petter S. Woll

**Author notes:** These authors contributed equally (Michael Hagemann-Jensen, Christoph Ziegenhain, Petter S. Woll).

## Abstract

Many biological processes are regulated by the direct interaction between two or more cell types. However, our understanding of the immediate dynamic changes in gene transcription upon physical interaction between two cells has remained limited due to technical limitations. Here we address these limitations in a model system of cancer-specific T cell receptor (TCR)-modified CD8 T cells where single and cancer-interacting T cells were isolated by image-enabled cell sorting and transcripts from heterotypic cancer cell and T cell pairs were *in silico* assigned. This approach uncovers immediate, dynamic changes in gene expression following the specific interaction between TCR-modified CD8 T cells and cancer cells. In addition to dissecting transcriptional cascades dependent on the peptide sensitivity of the TCR, we for the first time directly compare these gene expression changes between single T cells and T cells in direct physical contact with cancer cells. Modeling of the observed transcriptomic activation signature identifies phenotypically distinct tumor infiltrating CD8 T cell subsets associated with reduced TCR diversity in *in vivo* datasets. Taken together, the paradigm developed here allows for future clonal identification of T cell receptors mediating ongoing effective cytotoxic responses in vivo.

## Main text

Cellular interactions are critical for the correct regulation of biological systems throughout the human body^1, 2^. A prime example of this is the requirement for physical cell-cell interaction mediated by engagement of a specific T cell receptor (TCR) with cognate peptide presented on MHC class I expressed on a target cell for the activation of cytotoxic T cells. Although many signaling components in the cascade of protein-modifications downstream of TCR engagement have been previously identified^3^, gene expression changes and post-transcriptional regulatory mechanisms have often been explored at population rather than single level, and at timepoints where immediate and secondary effects are intermixed^4–9^. Single cell approaches have improved resolution towards uncovering transcriptional changes upon T cell activation^5, 6, 9–11^. However, it remains a challenge to reveal the initiating events at very early timepoints as these are often masked by the pool of pre-existing mRNA^12^. Single-cell studies also often lack information about the concrete cell pairing and the exact cellular composition of presumed interacting cells^5, 6, 9–11^. While spatial transcriptomics studies and sequencing of interacting cell doublets paired with statistical deconvolution of transcriptomes have provided some additional information on interacting cell pairs, they are restricted by low sensitivity and accuracy^4–6, 9^.

Here, we address these limitations with a novel methodical strategy where TCR specificity was controlled to allow comparison of TCR-specific and TCR-non-specific T cells. We here employed engineered primary T cells with TCRs targeting peptides of terminal deoxynucleotidyl transferase (TdT) or Melan-A (Mart1/control) presented on HLA-A*02:01, developed for TCR-guided immunotherapy against B/T-cell acute lymphoblastic leukemia (B/T-ALL)^13^ or melanoma^14^, respectively. The two TdT-specific TCRs have 4.8-fold difference in peptide sensitivity (T1_EC50_=5.8nM, T3_EC50_=1.2nM)^13^, resulting in enhanced killing of the TdT-expressing, Mart1-negative B-ALL cell line NALM-6 by T3 compared to T1 TCR T cells, and no effect of Mart1 TCR T cells (**Suppl. Fig. 1a**). To dissect the transcriptional changes leading to efficient killing, 1:1 co-cultures of TdT-expressing NALM-6 cells with T1, T3 or Mart1 TCR-modified T cells were subjected to image-enabled flow cytometry^15^ to precisely identify and isolate interacting T cell::NALM-6 pairs and single cells at different timepoints (**Fig. 1a; Suppl. Fig. 1b-h**; **Methods**). Interacting pairs were enriched by first gating for events staining positive for cell-surface markers exclusive to each cell type (CD19^+^CD3^+^CD8^+^mTCRb^+^). These interacting doublet populations were detected at higher frequency in the T1 and T3 compared to Mart1 co-cultures with NALM-6 cells (**Suppl. Fig. 1c**), in line with a recent study demonstrating that T cells with tumor-specific TCRs are enriched within interacting cell clusters obtained from clinical tumor isolates^11^, warranting focused analysis of T cells found in direct contact with other cell types. However, image-analysis of the CD19^+^CD3^+^CD8^+^mTCRb^+^ population revealed a heterogenous mixture of cells not only composed of a single T cell interacting with a single NAML-6 cell (**Suppl. Fig. 1d**), highlighting the importance of image-enabled gating for purification of true cell-cell pairs in direct contact (**Suppl. Fig. 1e-f**). Re-analysis of image-sorted T cell::NALM-6 pairs revealed that approximately 50% of the pairs remained in direct physical contact (**Suppl. Fig. 1g-h**), demonstrating that the interaction is strong enough to sustain the forces of cell sorting and reanalysis in a flow cytometer.

**Figure 1.**
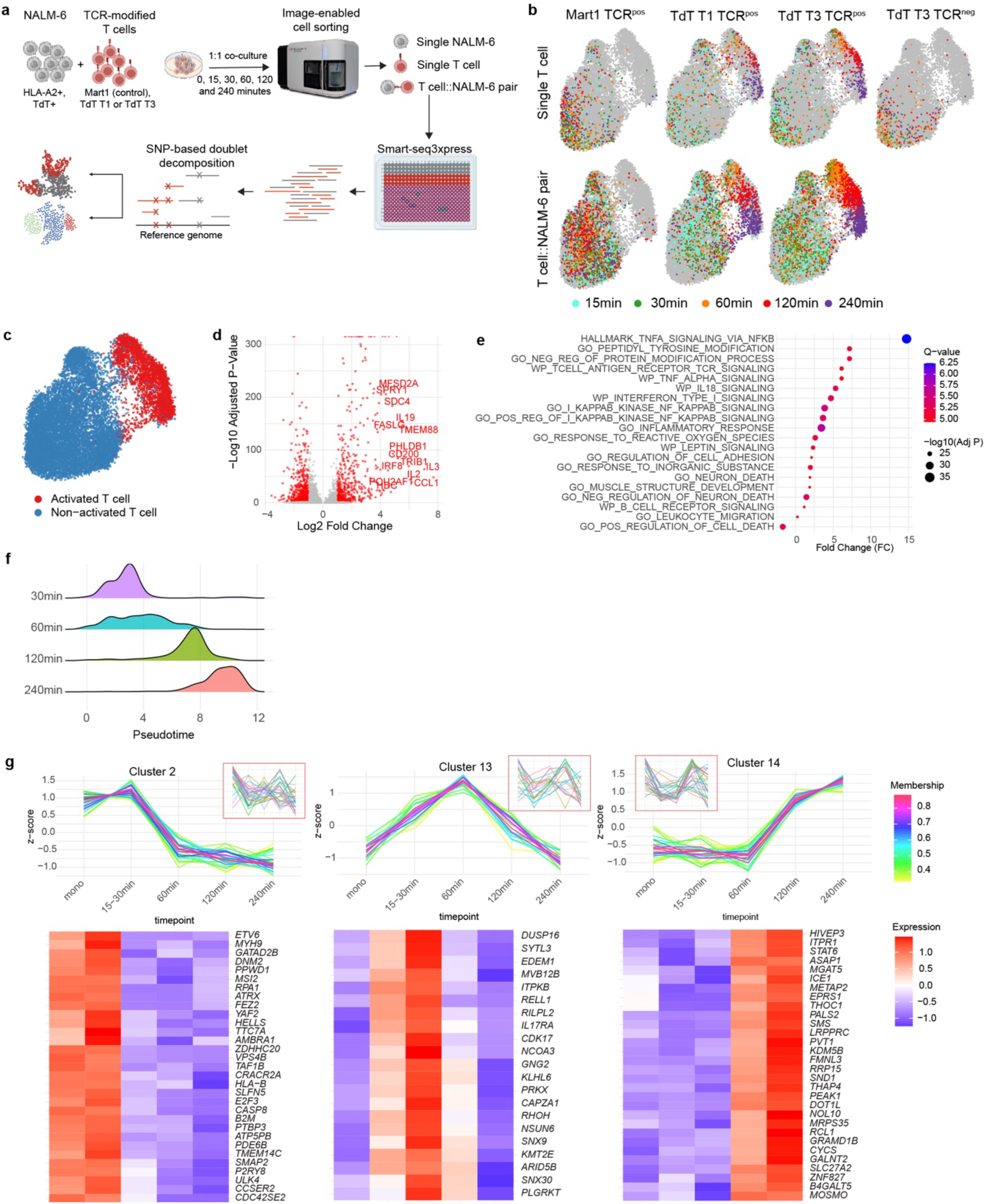
Decomposed unspliced pre-mRNAs uncover dynamic and transient transcriptional changes upon TCR-mediated activation. **a.** Overview of study design. NALM-6 cells were co-cultured with TCR-modified T cells (0-240 minutes) prior to image-enabled cell sorting that distinguished single NALM-6, single T-cell, and physically interacting T cell::NALM-6 pairs. Following Smart-seq3xpress RNA sequencing, single nucleotide polymorphisms (SNP) within transcript sequences were used to assign individual reads to the cell of origin. **b.** UMAPs of T cell specific gene expression levels from single T cells (top row) and T cell::NALM-6 pairs (bottom row) obtained at indicated timepoints (colored dots) following co-culture of NALM-6 cells with Mart1 (n=921 single and 2,611 pairs), T1 TdT (n=810 single and 2,214 pairs) or T3 TdT (n=1,281 single and 3,381 pairs) T cells. From T3 TdT co-cultures, both mTCRb positive (TCR^pos^, n=958) and mTCRb negative (TCR^neg^, n=323) single T cells were sorted, whereas from Mart1 and T1 only TCR^pos^ cels were analyzed. **c.** UMAP highlighting T cells (single and in direct contact with a NALM-6 cell) classified as transcriptionally activated (red) or non-activated (blue). **d.** Volcano plot highlighting differentially expressed genes (red indicating significance with adjusted p-value < 0.05 and absolute log2 fold-change > 1) in activated T cells compared to non-activated T cells (decomposed reads). **e.** Single cell pathway analysis of activated compared to non-activated T cells originating from single T cells and T cell::NALM-6 pairs (decomposed reads). **f.** Pseudotime plots of activated (combined single and paired T1 and T3 TCR CD8^+^ T cells) at different timepoints of culture. **g.** Representative temporal expression patterns (top row) and heatmap (bottom row) of gene clusters rapidly down-regulated (cluster 2), up-regulated early (cluster 13) or up-regulated late (cluster 14) in activated T1 and T3 cells following indicated minutes of co-culture. Shown are z-score normalized expression values (decomposed reads). Insert indicate the corresponding gene expression patterns observed in non-activated T cells. Additional patterns are displayed in **Supplementary** Figure 3.

Although T cells isolated from clinical tumor samples are enriched for tumor-specific T cells^11^, also non-specific T cells are found in direct contact with tumor cells (**Suppl. Fig. 1c**), requiring additional information to distinguish specific from non-specific TCR engagement. To assess the immediate transcriptional changes antigen-specific T cells undergo towards cytotoxic activation upon engagement with target cells, isolated T cell::NALM-6 pairs were subjected to full-length RNA-seq^16^ which allowed for two major advances. Firstly, we utilized genetic differences between NALM-6 and T cells to unambiguously assign transcripts containing single nucleotide polymorphisms (SNPs) to their cell of origin (false positive rate < 0.17%; **Suppl. Fig. 2a-e**). Furthermore, we focused the analysis on unspliced pre-mRNA molecules, which due to the rapid processing to spliced mRNA enriches for newly generated transcripts (**Fig. 1a, Methods**)^17^. Although this approach results in loss in read counts relative to the full dataset (**Suppl. Fig. 2f**), visualization of T cell specific reads using uniform manifold approximation and projection (UMAP) revealed an intermix of T cell::NALM-6 pairs and single T cells isolated after co-culture (**Suppl. Fig. 2g-h**), as also evident for single Mart1, T1 and T3 TCR T cells isolated prior to co-culture (**Suppl. Fig. 2i**). A clear time associated separation of a subset of the T1 and T3 TCR T cell transcriptomes was observed only in the TCR-modified (mTCRb^+^) T cells (**Fig. 1b**), and not in unspecific Mart1 TCR T cells, demonstrating a specific transcriptomic response upon specific TCR engagement. This unique time associated subset including only T1 and T3 mTCRb^+^ T cells was identified as transcriptionally activated T cells (**Fig. 1b-c**), enriched in a previously described gene expression signature associated with cytotoxic T cells having the capacity to execute serial killing (**Suppl. Fig. 2j**)^9^. Differential gene expression (DEG) analysis comparing activated and non-activated T cells identified 1,141 DEGs (**Fig. 1d**), enriched for several pathways associated with T cell activation^3^, including TCR, tumor necrosis factor alpha (TNFA), interferon (IFN) and nuclear factor kappa-light-chain-enhancer of activated B cells (NFKB) signaling pathways (**Fig. 1e**). This demonstrates that SNP-based decomposition of unspliced pre-mRNA from interacting T cell::NALM-6 pairs allows for accurate identification of transcriptional changes upon specific T cell activation.

As transcriptionally activated cells from different timepoints occupied distinct positions along pseudotime (**Fig. 1f**), we leveraged temporal information in the unspliced pre-mRNA to identify gene modules with dynamic expression during early T cell engagement. This revealed 16 temporal regulatory patterns (**Fig. 1g, Suppl. Fig. 3, Suppl. Table 1**). Focusing on transcription factors encoded by unspliced pre-mRNA transcripts we uncovered transient programs, including early induction of core NF-κB components (*REL, NFKB1*) followed by increased expression of NF-κB–responsive regulators such as *NFATC1, ZEB2*, and *STAT6* (**Suppl. Fig. 3**).

Interestingly, the effect on gene expression changes in interacting NALM-6 cells were much less pronounced than those observed in T cells, with only a subtle time associated effect following co-culture with both specific (T1 and T3) and non-specific (Mart1) TCR T cells (**Suppl. Fig. 4a-b**). A direct comparison of engaged NALM-6 cells with activated specific T cells to those in direct contact with non-activated specific T cells only revealed 8 DEGs (**Suppl. Fig. 4c**), of which 6 genes were shared DEGs detected in activated T cells (likely representing contaminating reads expected at the empirical false positive rate of 0.17%). This indicates that activation of antigen-specific T cells does not result in major target cell transcriptional changes in the 4-hour window investigated, in accordance with viability assays where the specific killing of NALM-6 cells in this time window is yet to occur (**Suppl. Fig. 4d-f**).

Next, we investigated to what extend the 10-fold higher efficiency in specific killing mediated by T3 compared to T1 TCR T cells (**Fig. 2a, Suppl. Fig. 1a**), is reflected in the transcriptional profiles of activated interacting T cells. Comparison between activated T1 and T3 cells at different timepoints revealed highly consistent gene expression (**Suppl. Fig. 5a**), indicating that the transcriptional response to target recognition is shared regardless of TCR peptide sensitivity. Both T1 and T3 TCR T cells had a significantly higher serial killer signature score compared to Mart1 cells after 60, 120 and 240 minutes of co-culture with NALM-6 cells (**Fig. 2b**). However, at each timepoint, T3 cells displayed significantly higher serial killer signature scores than T1 TCR T cells (**Fig. 2b**), alongside a greater fraction of transcriptionally activated cells (**Fig. 2c**) and a larger contribution to the leading edge of the molecular pseudotime trajectory (**Fig. 2d**). Taken together, our data suggest that the difference in peptide sensitivity primarily modulate the timing and fraction of T cells that become activated, while the transcriptional activation program itself is largely shared.

**Figure 2.**
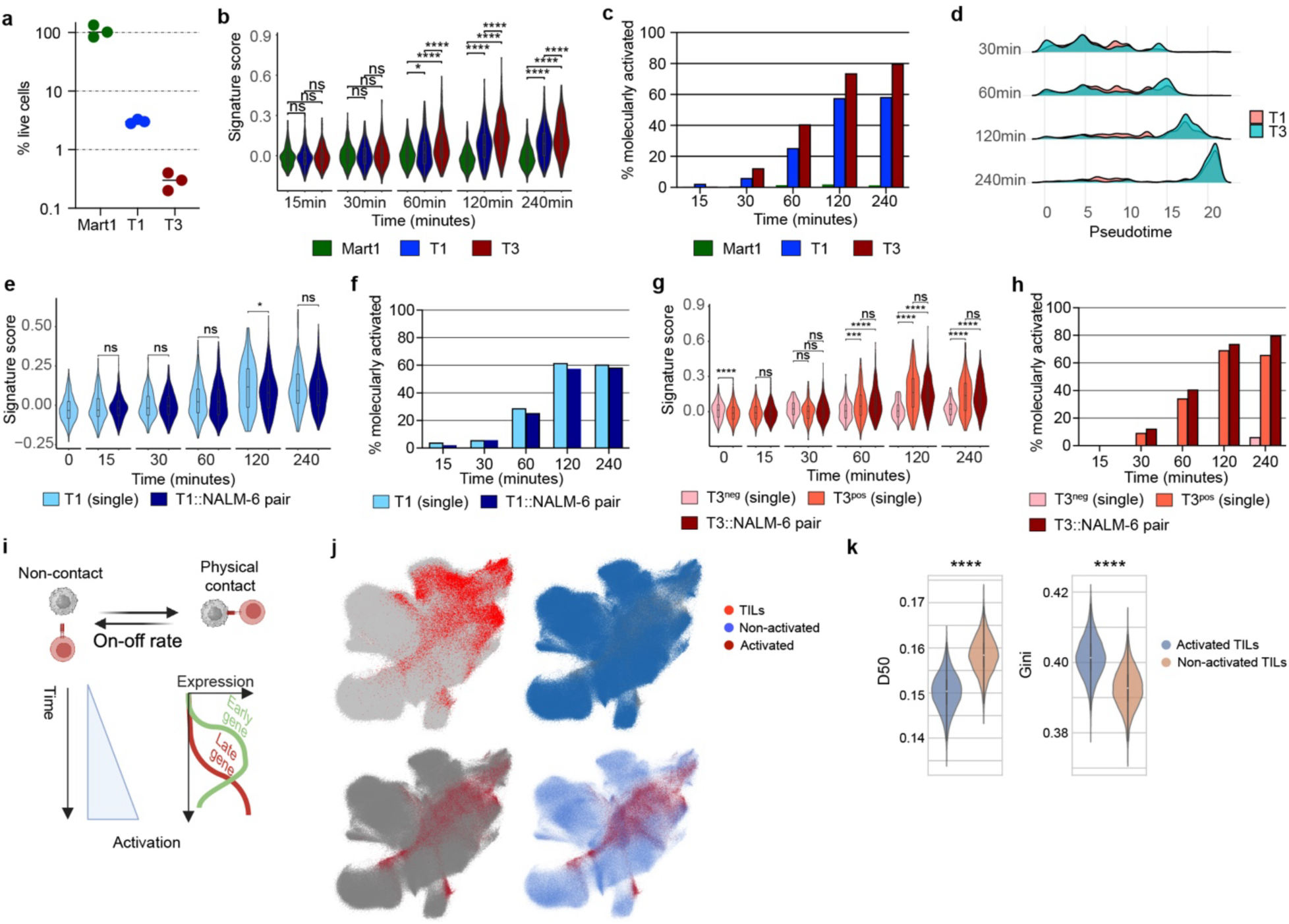
Coordinated transcriptional signatures in single and NALM-6 interacting T cells. **a.** Percentage live NALM-6 cells remaining following 48 hours *in vitro* killing assay with 1:1 co-culture with Mart1 (control), T1 and T3 TCR T cells (n=3). Numbers were compared to NALM-6 cell cultured for 48 hours in the absence of T cells. **b.** Violin plot of serial killer score within T cell::NALM-6 pairs at indicated timepoints of co-culture. Stars denote significance levels (* P<0.05; *** P<0.001; **** P<0.001; ns, not significant). **c.** Percentage of interacting T cell::NALM-6 pairs (Mart1, T1 and T3 T cells) present within transcriptionally activated cell clusters at indicated timepoints of co-culture. **d.** Pseudotime distribution of activated and non-activated T1 (red) and T3 (blue) TCR CD8^+^ T cells at different timepoints of culture. **e-f.** Violin plot of serial killer score (**e**) and percentage of total cells found within serial killer score positive cluster (**f**) of single T1 TCR T cells and T1 TCR cells in direct contact with a NALM-6 cell at indicated timepoints. **g-h.** Violin plot of serial killer score (**g**) and percentage of total cells found within serial killer score positive cluster (**h**) of single T3 TCR^neg^ or T3 TCR^pos^ T cells and T3 TCR^pos^ cells in direct contact with a NALM-6 cell at indicated timepoints. **i.** Model of successive T cell interactions with target cells leading to activation of T1 and T3 TdT TCR T cells. **j.** UMAP plots based on an existing CD8^+^ T cell atlas^19^ showing annotated tumor-infiltrating lymphocytes (TILs, bright red in top left panel), and cells following deep machine learning by Geneformer annotated as non-activated (blue, top right and bottom right) or activated (deep red, bottom panels). **k.** Violin plot of D50 (left) and Gini (right) index for TCR receptor diversity within activated and non-activated TILs.

Leveraging our SNP-decomposed transcriptomes, we were able for the first time to directly compare single T cells to single T cells in direct contact with a target cell. A large proportion of DEGs identified from SNP-decomposed reads (**Fig. 1d)** were shared with DEGs identified when performing the same analysis on reads obtained from single T cells without SNP-decomposition (**Suppl. Fig. 5b-c**). Overall, both single T cells and T cell::NALM-6 pairs, independent TCR sensitivity, showed high gene expression correlation, similar serial killer score and contribution to the transcriptionally activated cell cluster (**Fig. 2e-h, Suppl. Fig. 5d**). Inclusion of T3 TCR-negative T cells from the same co-culture showed no increase in serial killer score or contribution to the transcriptionally activated cell clusters over time (**Fig. 2g-h**), revealing that activation of the T1 and T3 single cells is specifically interaction-triggered and not driven by bystander effects in the co-culture. Thus, our data provides a model where a shared transcriptional activation of specific T cells is driven by successive physical interactions between the TCR-specific T cells and target cells, generating a transcriptional program that eventually leads to robust killing of the NALM-6 cells (**Fig. 2i**).

To demonstrate how our early, dynamic transcriptional program can be used to identify candidate *in vivo* cytotoxic effector cells, we fine-tuned a large single-cell foundation model^18^ (pre-trained on >100M single-cell transcriptomes) using gene expression profiles from activated T1/T3 TCR^+^ CD8^+^ T cells. AI-guided predictions performed on an atlas of >1 million CD8⁺ T cells from different tissue sites and disease conditions, including a large subset of tumor-infiltrating lymphocytes (TILs)^19^, classified 102,745 CD8^+^ T cells as transcriptionally activated (**Fig. 2j**). Supporting the specificity of the activation gene signature, naïve, central memory and terminal differentiated effector memory T cell subsets had low (<2.9%) content of predicted activated cells (**Suppl. Fig. 6a**). Intriguingly, high fractions of transcriptionally activated T cells were found in multiple T cell subtypes (**Suppl. Fig. 6a**), including a significant enrichment of activated cells among TILs (25.07% contribution, odds ratio 5.33, p-value <0.0001, **Fig. 2j**). This demonstrated that conventional gene expression clustering and previously defined gene signatures (**Suppl. Fig 6b**) do not capture the cell populations predicted by the expression signature defined in this work. We next explored the functional implications within distinct T cell subsets by comparing the TCR diversity of the transcriptionally activated cells. In TILs (**Fig. 2k**) and in the T cell subsets with the highest proportion of transcriptionally activated cells (ITGAE^+^ T_ex_, a cytotoxic subset enriched in solid tumors and associated with good response to PD-L1-immune checkpoint therapy^19, 20^, and CREM^+^ T_rm,_ enriched in TILs^19^) (**Suppl. Fig 6c**), transcriptionally activated cells had a significantly lower TCR diversity compared non-activated T cells. In combination with reliable TCR sequence reconstruction^16^, our approach therefore supports identification and clonotypic characterization of T cell subsets with an activated signature also from clinical samples.

Together, our findings reveal an immediate and dynamic transcriptional signature that governs productive T cell activation following transient, physically validated target-cell engagement. By combining precise image-enabled isolation of true interacting cell pairs with SNP-decomposed full-length single-cell transcriptomes, we dissect the immediate gene-expression events that precede cytotoxic function, independently of pre-existing mRNA or bystander effects. Despite differences in peptide sensitivity, T1 and T3 TCR T cells were activated through a shared kinetic expression program, with higher sensitivity primarily increasing the frequency of activated cells rather than altering the activation pathway architecture. Finally, by fine-tuning a large-scale single-cell foundation model, we demonstrate that this early activation signature is robust, specific, and broadly applicable for identifying poised cytotoxic T cells across diverse inflammatory and tumor microenvironments, underscoring its translational potential for immunotherapy discovery and precision T-cell engineering.

## Methods

### Cell lines and generation of TCR-modified T cells

NALM-6 cells obtained from Deutsche Sammlung von Mikroorganismen und Zellkulturen were maintained in RPMI medium (ThermoFisher) supplemented with 10% fetal calf serum (FCS; ThermoFisher) and 1% penicillin/streptomycin (P/S; PAA Laboratories). T cells expressing TdT-specific or Mart1-specific (DMF5) T cell receptors (TCRs) were generated as previously described^13, 14, 21^. Briefly, peripheral blood mononuclear cells from an HLA-A*02 positive healthy donor were transduced using retroviral constructs, expanded and frozen down for later experiments. Vials containing T1 TdT, T3 TdT or Mart1 TCR-modified cells, all originating from the same healthy donor were thawed 5-6 days prior to setting up the co-culture. The thawed T cells were maintained in X-vivo-20 medium (Lonza) supplemented with 5% human serum (Trina Biotech), 1% P/S, 5ng/mL IL-7 (PeproTech) and 5ng/mL IL-15 (PeproTech). Two separate experiments were performed where the T cells originated from the same healthy donor but represented two different TCR transductions, where experiment 1 included TdT T3 and Mart1 from the same transduction and experiment 2 included TdT T1, TdT T3 and Mart1 from the same transduction. The transduction efficiency, evaluated by staining for the mouse TCRβ chain (mTCRb)^13^, was between 65-70%.

### *In vitro* co-culture of T cells and target cells

NALM-6 and T cells were harvested and counted on a hemacytometer with Tryphan blue dye exclusion for dead cells. Co-cultures were initiated by combining 100,000 NALM-6 cells with 100,000 T cells (1:1 ratio) in individual wells of a U-shaped 96-well plate in 200μL X-vivo-20 medium supplemented with 5% human serum. Monocultures, representing the baseline (0) timepoint, were composed of only NALM-6 or TCR-modified T cells. Co-cultures were incubated at 5% CO2 and 37°C for 15, 30, 60, 120 or 240 minutes for RNA sequencing or 48 hours for evaluation of *in vitro* killing efficiency. After incubation, each well was harvested by gentle pipetting, transferred to 1.5mL Eppendorf tubes, washed with ice-cold phosphate-buffered saline without magnesium and calcium (DPBS; Gibco) supplemented with 5% FCS and prepared for flow cytometry analysis.

### *In vitro* killing analysis

For quantification of *in vitro* killing, 100,000 NALM-6 cells were co-cultured with 100,000 TdT T1, TdT3 or Mart1 T cells in a 96-well round-bottom plate in triplicates. Cells were washed and stained after 48 hours of culture with the following antibodies: CD8a-APC-Cy7 (clone RPA-T8, BioLegend), HLA-A2-BV650 (clone BB7.2, BioLegend), CD19-BV785 (clone HIB19, BioLegend), CD3-BB515 (clone HIT3a, BD), CD10-PECy5 (clone HI10a, BioLegend) and CD4-PECy7 (clone RPA-T4, BioLegend). After 15 minutes of staining on ice, the cells were washed and resuspended in DPBS with 5% FCS and 10,000 CountBright Absolute Counting Beads (ThermoFisher). 4′,6-diamidino-2-phenylindole (DAPI; ThermoFisher) was used for exclusion of dead cells. The samples were acquired on a BD LSRFortessa where 3,500 beads were recorded for each well. The percentage live cells were calculated in relation to NALM-6 cells in monocultures.

### Image-enabled flow cytometry cell analysis and sorting

Harvested washed cells were spun at 400g for 5 min and incubated with human Fc block reagent (Miltenyi Biotec) for at least 10 minutes at 4°C before staining for 15 minutes on ice with the following antibodies: CD3-BB515 (clone HIT3a, BD), CD19-PECy7 (clone HIB19, BD)/CD19- RB780 (clone HIB19, BD), mouse TCRβ-PE (clone H57-597, BioLegend)/mouse TCRβ-RB613 (clone H57-597, BD) and CD8a-APC -Cy7 (clone RPA-T8, BioLegend). DAPI was included for detection of dead cells. Acquisition was performed on a BD FACSDiscover™ S8 (BD)^15^ with a 100μm nozzle, available within the MedH Flow Cytometry Core facility at the Karolinska Institutet. As CD19 and CD3 represented mutually exclusive markers for NALM-6 and T cells, respectively, these were used to enrich for interacting NALM-6 and T cells (double positive), as well as identifying individual NALM-6 and T cells (single positive) (**Suppl. Fig. 1b**). Image-enabled parameters including delta center of mass and radial moment for CD3, CD19 and mouse TCRβ was used to purify cells as single cell or in direct physical interaction (**Suppl. Fig. 1e-f**). Prior to sorting accurate deposition into 384-well plates was confirmed by sorting co-cultured cells labelled with 250μg/mL horseradish peroxidase (ThermoFisher) into 3μL TMB High Sensitivity substrate solution (BioLegend), resulting in blue color-conversion in >98% of the wells. Index sort information and manual investigation of images using the CellView plugin (version 2.0.15) in FlowJo (version 10.10.0) was used to annotate each sorted cell to allow exclusion of cells not in direct contact, cells with abnormal morphology or cells with ambiguous phenotype (e.g. unable to confidently identify cell as T cell or NALM-6 cell).

### RNA library generation and sequencing

Single cells and doublets were sorted into 384-well plates and prepared for Smart-seq3xpress library preparation as previously described^16, 22^. In brief, single cells and doublets were sorted into lysis buffer and amplified using 15 cycles of PCR after cDNA conversion. 1μL of the diluted amplified cDNA was subsequently tagmented using 0.007μL TDE1 (Illumina) per well, before index PCR with Illumina Nextera index primers for 14 cycles. Afterwards libraries were pooled, bead-cleaned and evaluated via Qubit (Thermo Scientific) and Bioanalyzer (Agilent). Single-cell RNA sequencing libraries were sequenced on MGI DNBSEQ G400 and DNBSEQ T7 platforms using universally compatible (App-D) sequencing reagents according to the manufacturer’s instructions.

### Read alignment and transcript quantification

Raw FASTQ files were processed using zUMIs (v2.9.7)^23^. Two batches of data were processed separately. In the YAML configuration, batch 1 read 1 was defined with find_pattern: ATTGCGCAATG and base_definition: UMI(12–21); cDNA(25–150). Batch 1 read 2 was parsed with cDNA (1–150) and barcode (BC, 1–20), while batch 2 read 2 was parsed with cDNA (1–100) and BC (101–120). UMIs were collapsed using a Hamming distance of 1, and barcode binning was set to 1. Reads were aligned to the human reference genome (hg38) using STAR^24^ (v2.7.3a) with gene annotations from GENCODE GRCh38.p14 (Release 47). Additional STAR parameters included --clip3pAdapterSeq CTGTCTCTTATACACATCT and --limitOutSJcollapsed 3000000. Gene-level read and UMI count tables were generated, capturing counts from exons, introns, and overlapping regions. To more enhance quantification of actively transcribed genes rather than the steady-state levels of mature mRNA, only the reads aligned to annotated introns and intron-exon overlapping regions representing unspliced pre-mRNA were used for all downstream analysis.

### Variant calling and cell-specific read separation

VCID (Variant-based Cell-cell Interaction Deconvoluting) was developed to perform read mapping, SNP calling, differential SNP filtering and formatting, and read separation starting from RNA-seq FASTQ files. BAM files were first processed with bamaddrg to assign identifiers to each cell type, including T single cells, NALM-6 single cells, and doublets. Single nucleotide variants (SNVs) were detected using FreeBayes^25^ (v1.3.8) with the parameters --min-mapping-quality 10, --min-base-quality 20, and --min-alternate-count 5 to generate variant call format (VCF) files. VCF files were further filtered and formatted using BCFtools^26^. Only variants of type SNP were retained using bcftools filter, and genotypes containing the alternative allele 2 were excluded, keeping only genotypes represented as 0/0, 0/1, or 1/1. Cell-specific reads were then separated based on informative SNPs, generating cell-specific BAM files and corresponding gene-level expression count matrices.

### Single-cell clustering and quality control

For data before read separation, dimensionality reduction and clustering were performed using Seurat (v5.3.0)^27^. Cells were filtered using the criteria pct_coding*100 ≥ 50, nreadpairs > 10000, and mitoFrac < 0.3, and only genes expressed in at least five cells were retained, resulting in 18,875 cells. Data were normalized using NormalizeData with method “LogNormalize” and a scale factor of 10,000. Cell cycle scores were calculated using CellCycleScoring with S-phase genes (s.features) and G2/M-phase genes (g2m.features) from cc.genes.updated.2019.

Highly variable genes were identified with FindVariableFeatures using the “dispersion” method, selecting 2,000 genes. Data scaling was performed with ScaleData, regressing out nCount_RNA, S.Score, and G2M.Score while scaling only the highly variable genes. Batch effects were corrected using RunHarmony^28^ with “batch” as the covariate. Harmony-corrected embeddings were used for FindNeighbors (dims = 1:30) and clustering with FindClusters using the SLM (Smart Local Moving) algorithm at a resolution of 0.8. Two clusters showing extremely low nCount_RNA were removed, resulting in 18,009 cells and 26,738 features. For data after read separation, similar procedures were applied with adjusted cell quality filtering criteria. For T cell-specific reads, cells were filtered with pct_coding*100 ≥ 50, nreadpairs > 200, mitoFrac < 0.3, and nGenes_in_Tcell > 150, yielding 13,919 cells and 18,301 features. For NALM-6-specific reads, cells were filtered with pct_coding*100 ≥ 50, nreadpairs > 200, mitoFrac < 0.3, and nGenes_inex_Nalm6cell > 600, resulting in 12,000 cells and 17,388 features.

### Differential gene expression and pathway analysis

Differentially expressed genes were identified using FindMarkers in Seurat with the Wilcoxon rank-sum test. For the T cell–specific expression data after read separation, three clusters with a resolution of 0.8 were defined as activated T cells (activate_state == “act”), while all remaining clusters were defined as non-activated T cells (activate_state == “nonact”). SCPA^29^ was then applied to compare activated T cells and non-activated T cells to identify pathways enriched during T cell activation. Differentially expressed genes were identified based on log2-fold change >1 or <-1 and adjusted p-value <0.05.

### Clustering of gene expression patterns during T cell activation

After read separation, the T cell-specific log-normalized expression matrix was used for gene pattern analysis with TCseq^30^. Genes were first filtered based on differential expression, retaining those with p < 0.05 and absolute log fold change > 1. Then expression differences between adjacent time points were calculated, and genes with at least one difference > 0.25 were selected, resulting in 1,104 candidate genes for clustering. Cells at 0 min were defined as non-cultured T cells with activate_state == “nonact”. Cells at 15, 30, 60, 120, and 240 min were defined as T1 and T3 cells with activate_state == “act”. As a baseline control, non-cultured nonact T cells were used as 0 min, and T1 and T3 cells with activate_state == “nonact” were used for subsequent time points.

### Deep learning–based activation state classification

Geneformer^18^ (v2) was applied to T cell-specific data after read separation. Cells previously clustered at a resolution of 0.6 were manually annotated into two major states, resting state and activated state. The Geneformer Classifier function was then used to fine-tune the model with these annotated expression profiles, enabling discrimination between activated and non-activated T cells. The fine-tuned classifier was subsequently applied to an external T cell dataset^19^ to infer activation states.

## Code availability

Procedures for read mapping, SNP calling, read separation, and gene/UMI counting has been integrated into VCID (Variant-based Cell-cell Interaction Deconvoluting) and the code utilized for the analysis is available on GitHub at: https://github.com/ziegenhain-lab/VCID

## Acknowledgements

We thank the MedH Flow Cytometry Core Facility financed by the Infrastructure Board at Karolinska Institutet for providing instruments and assistance for cell sorting. This work was supported by funding from Cancerfonden (22 2178 Pj to P.S.W.), Radiumhemmets Forskningsfonder (244282 to P.S.W.), Karolinska Institutet (P.S.W.), Åke Wiberg foundation (M22-0019 & M25-0084 to C.Z., M22-0151 to M.H-J.), the Swedish Research Council (2022-01471 to C.Z., 2023-02564 to M.H-J.), the Frontier Grant of the Department of Medical Biochemistry and Biophysics (Karolinska Institutet, to C.Z.), KI Foundation grant (2024-02674 to M.H-J.), Jeanssons Stiftelser grant (J2023-0034 to M.H-J.) and Swedish Society for Medical Research (SSMF, SG-25-0250-B to M.H-J.).

## Contributions

M.H-J., C.H. and P.S.W. conceived and designed the study. Z.Q., M.L, N.S. and E.G. performed experiments. Z.Q. performed and interpreted data analysis with input from M.H-J., C.H. and P.S.W. Z.Q., M.H-J., C.H. and P.S.W. generated figures and wrote the manuscript. All authors have read and approved the manuscript.

---

**Supplemental Figure 1.**
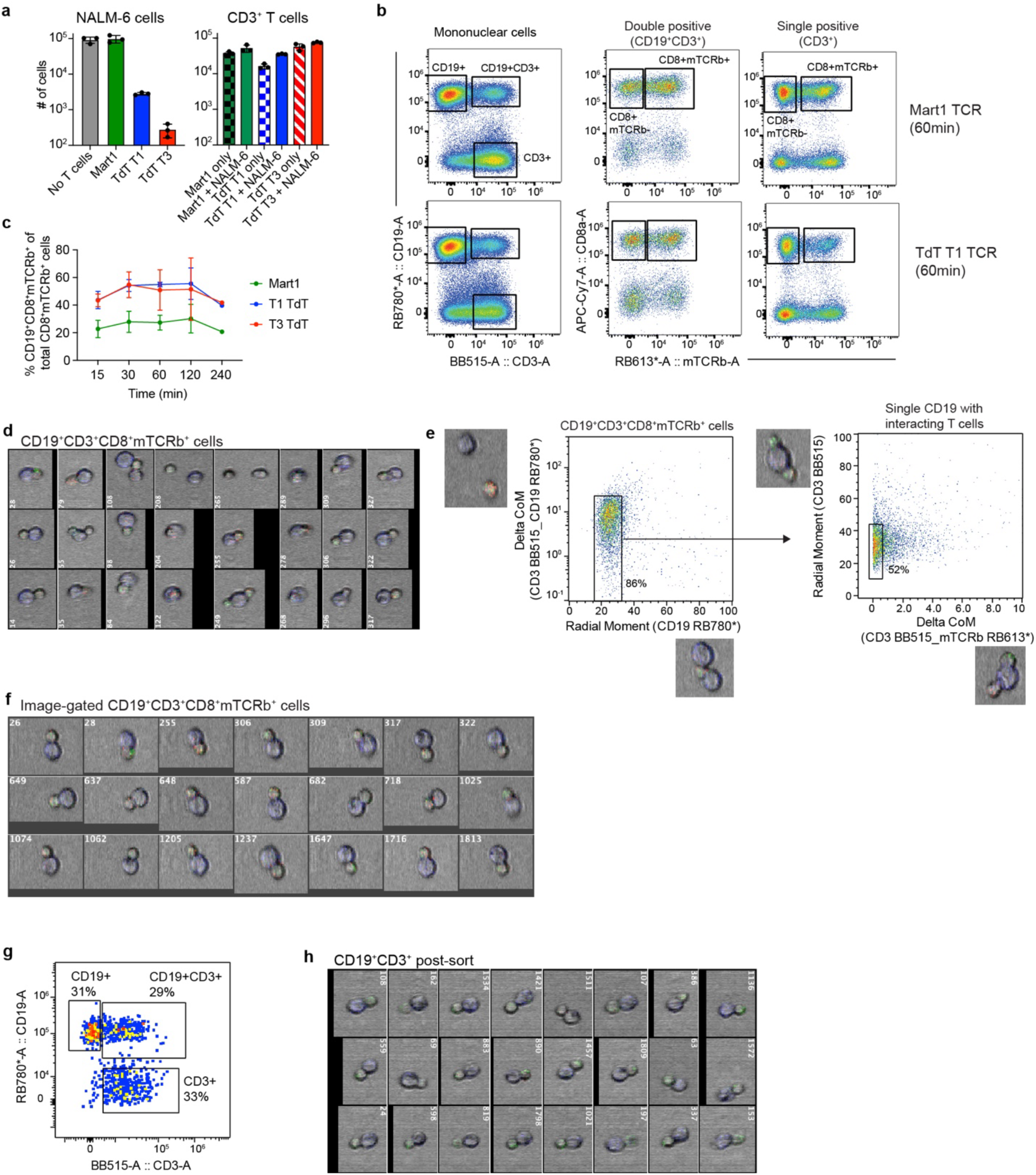
Image-enabled cell sorting allows purification of physically interacting T cell::NALM-6 pairs. **a.** Mean (SEM) number of NALM-6 and TCR-modified (Mart1, TdT T1 or TdT T3) CD3^+^ T cells remaining following 48 hours 1:1 co-culture (n=3 for all conditions). **b.** Representative gating strategy of Mart1 (top row) and T1 TdT (bottom row) TCR T cells co-cultured with NALM-6 cells for 60 minutes allowing enrichment of interacting T cells and NALM-6 cells based on expression of the mutually exclusive cell surface markers CD19 (NALM-6 specific) and CD3 (T cell specific). **c.** Mean (SEM) percentage of CD8^+^mTCRb^+^ cells found within CD19^+^CD3^+^ gate compared to the total CD8^+^mTCRb^+^ cells detected in the well (n=3) at indicated timepoints of co-culture. **d.** Images of CD19^+^CD3^+^CD8^+^mTCRb^+^ cells without image-based gating showing a heterogeneous pattern, including multiple NALM-6/T cells in direct contact with a T-/NALM-6 cell (#55, 108, 249, 327), cell without physical contact (#208, 265) and ambiguous cellular interaction (#122, #204, 268, 278), in addition to a single NALM-6 cell physically interacting with a single T cell. **e-f.** Representative FACS plots (**e**) and images (**f**) showing image-enabled gating strategy to remove non-interacting cell pairs and multiple NALM-6 or T cells, to recover a purified population composed of a single NALM-6 cell in direct contact with a single T cell. Percentages indicate cells within each gate based on the parental population. **g-h.** Post-sort analysis of T cell::NALM-6 pairs sorted based on image-enabled gating as shown in **e**. Re-analysis of sorted cells shows that approximately 50% of sorted T cell::NALM-6 pairs remain in direct contact as indicated by co-expression of CD19 and CD3 (**g**) and analysis of images from the recovered cells (**h**).

**Supplemental Figure 2.**
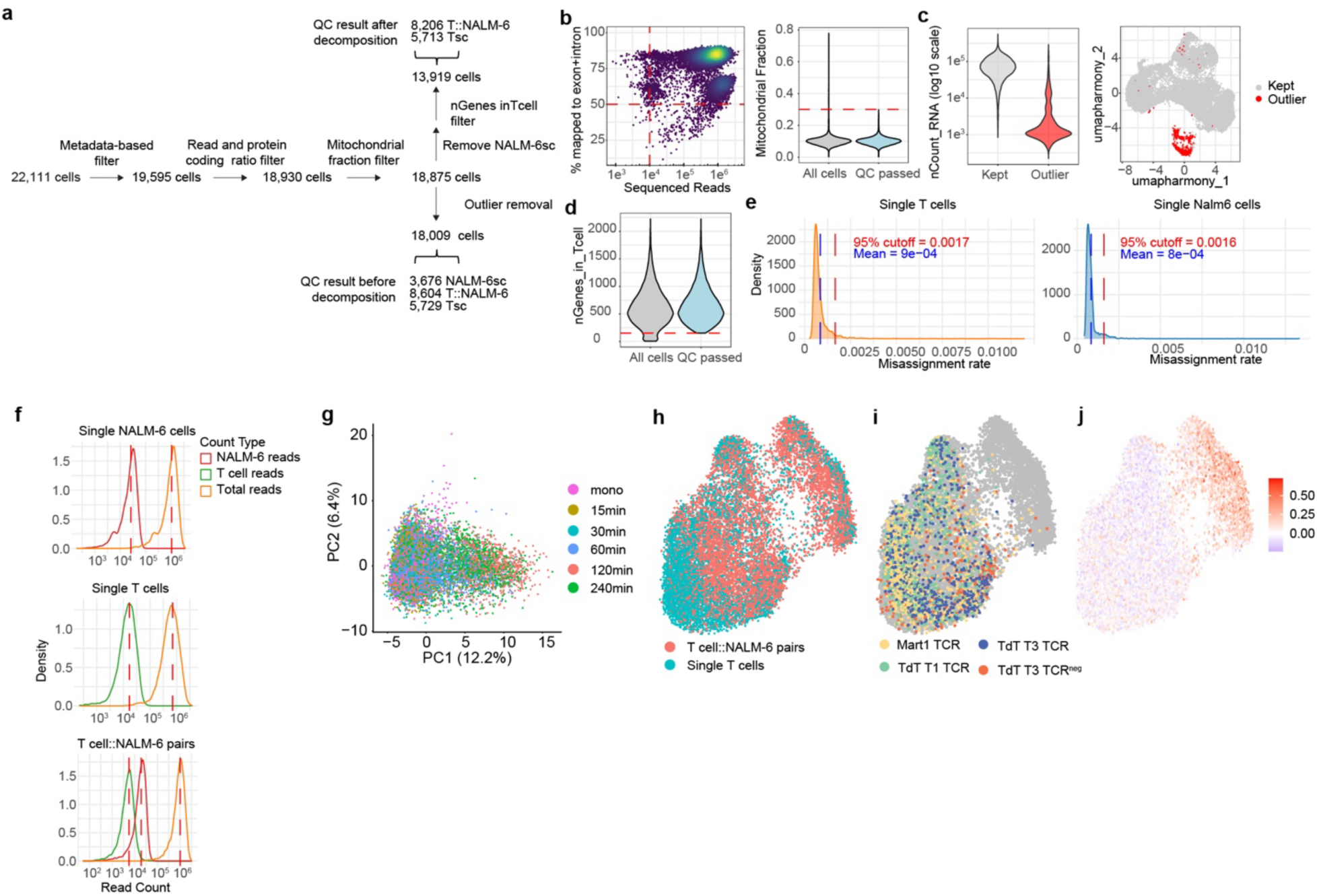
SNP-based read decomposition allows transcriptional comparison of single cells to T cell::NALM-6 pairs. **a.** Flow chart overview of cells included in the analysis after applying quality control filtering. Cells remaining after quality control (QC) before and after read decomposition are shown for single NALM-6 cells (NALM-6sc), single T cells (Tsc) and T cell::NALM-6 pairs. Metadata based filter was based on visual inspection of images of sorted single cells of cell pairs to remove cells not in direct contact and apoptotic/dead cells based on DAPI staining. **b.** 2D distribution of read depth and percentage mapping to exon or intronic regions (left plot) and fraction mitochondrial RNA (right plot) in cells before read decomposition. Red lines indicate cut-off for inclusion in analysis. **c.** Violin plot showing nCount RNA distribution (left) and UMAP plot (right) for outliers excluded from the analysis (red) and cells included in the analysis (grey) based on reads before applying read decomposition. **d.** Violin plot showing distribution of genes detected as T cell-specific in cells after read decomposition. Cells above the red line (>150 genes) were included in the analysis. **e.** Density plot showing misassignment rate of reads from single T cells (left) and single NALM-6 cells (right) after read decomposition. Blue lines indicate mean and red lines indicate 95% cut-off. **f.** Density plot showing read count distribution in single NALM-6 (top), single T cells (middle) and T cell::NALM-6 pairs (bottom). Red and green lines indicate intronic reads assigned to NALM-6 cells and T cells, respectively, after read decomposition, whereas orange lines indicate total intronic reads before read decomposition. **g.** Principal component analysis (PCA) plot based on SNP-decomposed T-cell specific reads from indicated timepoints. Percent variance indicated within parenthesis. **h-j.** UMAPs based on SNP-decomposed T cell specific reads highlighting single or paired configuration (**h**), uncultured (0 min) T cells (**i**), and heatmap of serial killer score (**j**).

**Supplemental Figure 3.**
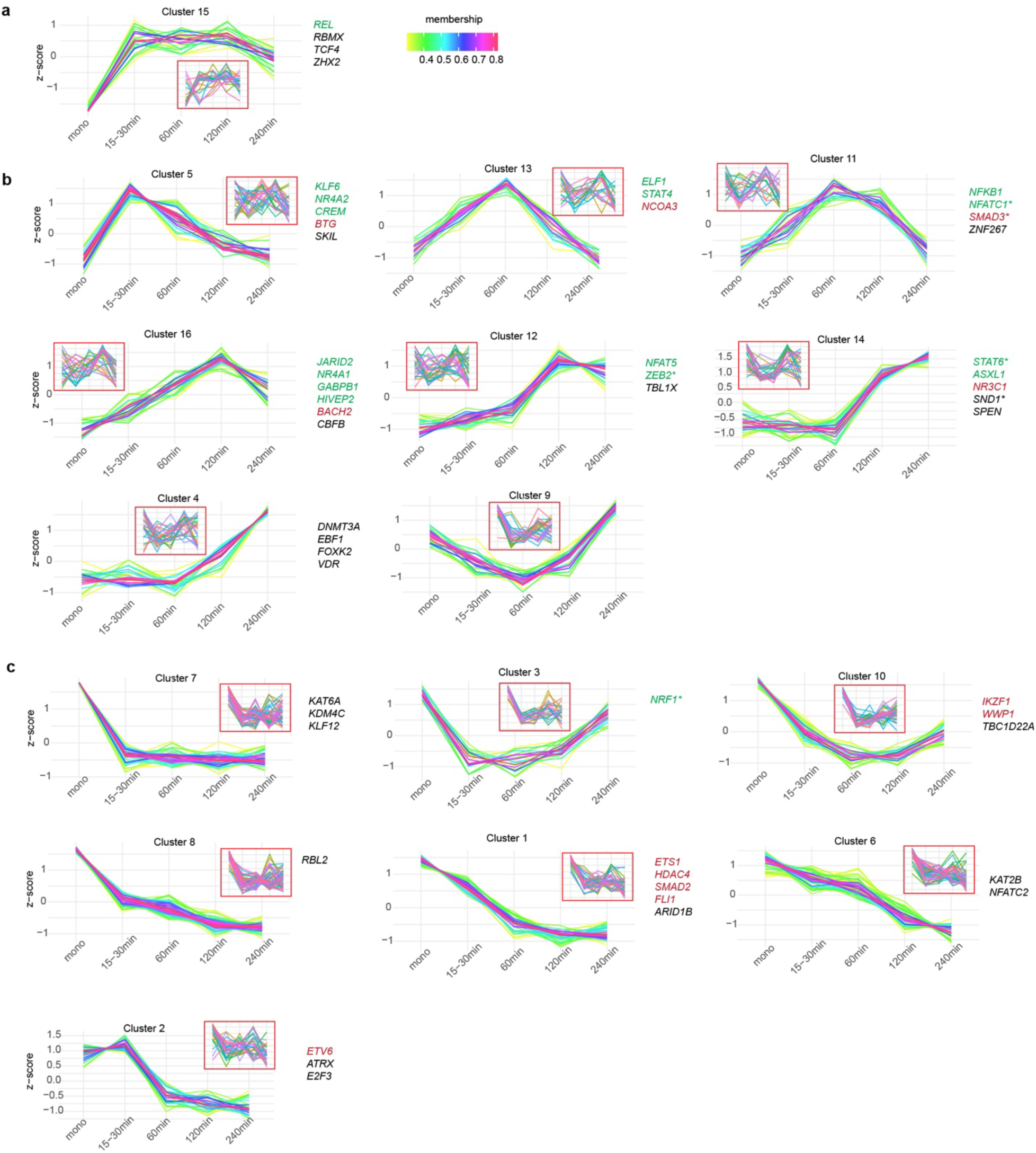
Temporal gene expression changes in activated single T cells and T cells in direct contact with NALM-6 cells. **a-c.** Expression patterns of stable (**a**) or transient (**b**) up-regulated and down-regulated (**c**) gene expression clusters identified in activated T1 and T3 cells following indicated minutes of co-culture. Expression values were z-score standardized and based on the read-separated quantification. Insert indicate the corresponding gene expression patterns observed in non-activated T cells. Annotated transcription factors highlighted based on association with T cell activation (green font) or suppression (red font) are indicated to the right of each plot. All genes associated with each gene cluster are shown in **Supplemental Table 1**.

**Supplemental Figure 4.**
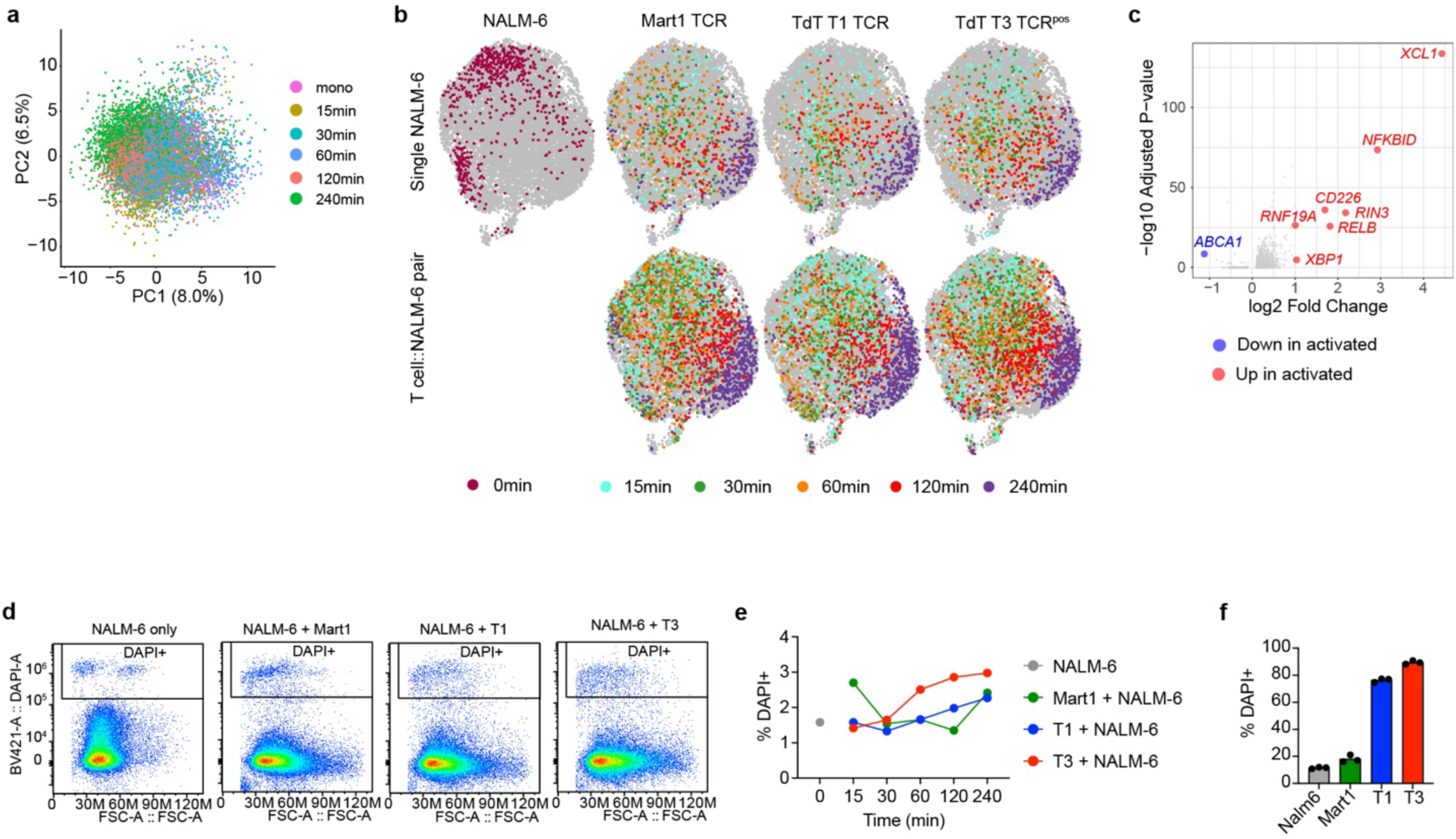
Limited immediate transcriptional impact in NALM-6 cells paired with activated T cells. **a.** PCA plot based on SNP-decomposed NALM-6 specific reads from indicated timepoints. Percent variance indicated within parenthesis. **b.** UMAPs based on SNP-decomposed NALM-6 specific reads in single NALM-6 cells (top row) and T cell::NALM-6 pairs (bottom row) obtained from indicated timepoints (colored dots) following co-culture of NALM-6 cells with Mart1 (n=909 single and 2,651 pairs), T1 TdT (n=833 single and 2,298 pairs), T3 TdT (n=1,270 single and 3,389 pairs). **c.** Volcano plot highlighting differentially expressed genes (red/blue indicate significance with adjusted p-value < 0.05 and absolute log2 fold-change >/<1) in NALM-6 cells paired with an activated T cell compared to NALM-6 cells paired with a non-activated T cell (decomposed reads). **d-e.** Representative FACS plots (d) showing percentage DAPI positive cells within CD19^+^ NALM-6 cells before co-culture and after 240 min co-culture with Mart1, T1 and T3 TCR T cells. Quantification of DAPI positive cells (e) within CD19^+^ NALM-6 cells at indicated timepoints of co-culture with TCR T cells. **f.** Quantification of DAPI positive cells within CD19^+^ NALM-6 cells after 48 hours of culture with or without TCR T cells.

**Supplemental Figure 5.**
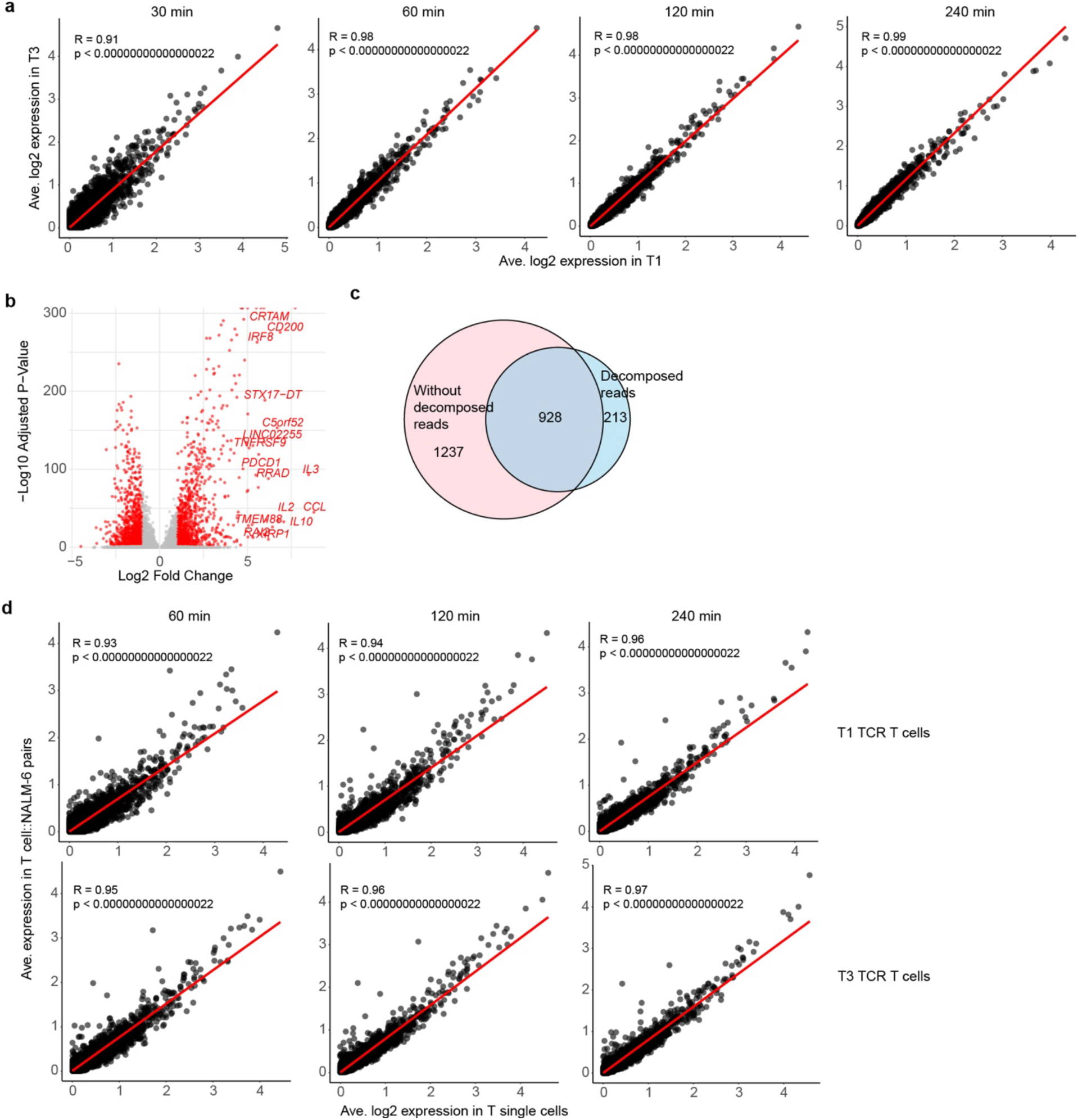
Gene expression of transcriptionally activated cells is not dependent on TCR strength or cellular configuration. **a.** Correlation in gene expression between T1 (x-axis) and T3 (y-axis) TdT TCR T cells at 30, 60, 120 and 240 min incubation. Correlation co-efficient (R) and p-value is shown. **b.** Volcano plot highlighting down- (left side) and up-regulated (right side) genes in activated T cells compared to non-activated T cells before read decomposition. Red indicates significance with adjusted p-value < 0.05 and absolute log2 fold-change >/<1. Cells were assigned as activated and non-activated based on contribution to clusters marked by high serial killer score. **c.** Venn diagram showing overlap between differentially expressed genes (adjusted p-value < 0.05 and absolute log2 fold-change >/<1) between activated and non-activated T cells before and after read decomposition. **d.** Correlation in gene expression between T1 (top row) or T3 (bottom row) TdT TCR T cells in single cell (x-axis) or T cell::NALM-6 pair (y-axis) configuration.

**Supplemental Figure 6.**
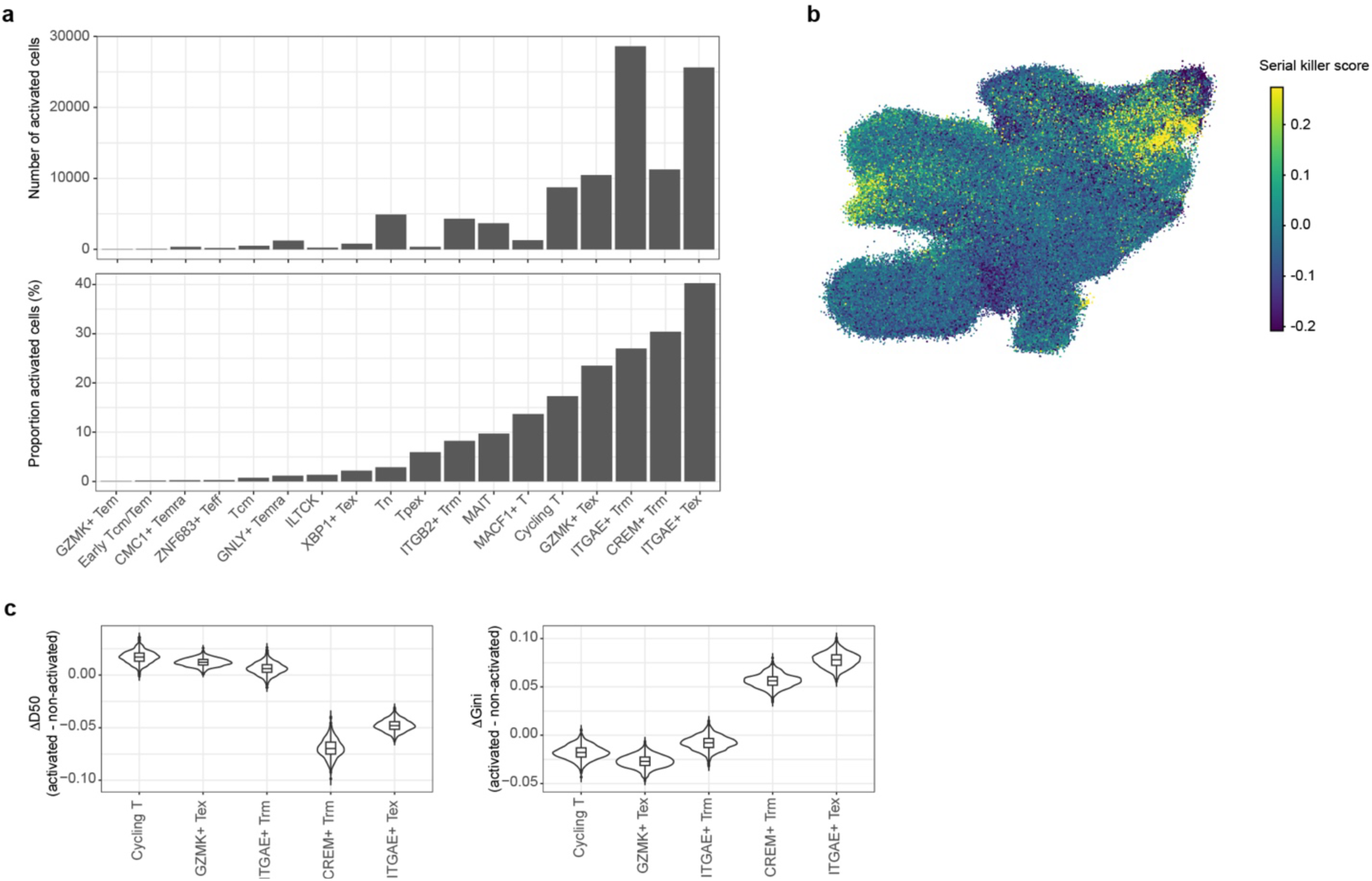
Gene signature for transcriptionally activated CD8^+^ T cells enriches for distinct T cell subsets with lower TCR diversity. **a.** Number of cells (top panel) and proportion of cells (lower panel) within indicated CD8^+^ T cell subsets classified as transcriptionally activated following deep machine learning by Geneformer. Annotation was derived from original data set by Xue et al^19^. **b.** UMAP heatmap of serial killer score within an existing CD8^+^ T cell atlas by Xue et al^19^. **c.** Difference (Δ) in D50 (left) and Gini (right) index for TCR repertoire diversity in the top 5 T cell subsets with the highest proportion of activated cells. The Δ was calculated by subtracting the non-activated cell index from the activated cell index.

